# *Garcinia kola* and garcinoic acid suppress SARS-CoV-2 spike glycoprotein S1-induced hyper-inflammation in human PBMCs through inhibition of NF-κB activation

**DOI:** 10.1101/2021.05.18.444690

**Authors:** Olumayokun A Olajide, Victoria U Iwuanyanwu, Izabela Lepiarz-Raba, Alaa A Al-Hindawi, Mutalib A Aderogba, Hazel L Sharp, Robert J Nash

## Abstract

Symptoms and complications associated with severe SARS-CoV-2 infection such as acute respiratory distress syndrome (ARDS) and organ damage have been linked to SARS-CoV-2 spike glycoprotein S1-induced increased production of pro-inflammatory cytokines by immune cells. In this study, the effects of an extract of *Garcinia kola* seeds and garcinoic acid were investigated in SARS-CoV-2 spike glycoprotein S1-stimulated human PBMCs. Results of ELISA experiments revealed that *Garcinia kola* extract (6.25, 12.5 and 25 μg/mL) and garcinoic acid (1.25, 2.5 and 5 μM) significantly reduced SARS-CoV-2 spike glycoprotein S1-induced increased secretion of TNFα, IL-6, IL-1β and IL-8 in PBMCs. In-cell western assays showed that pre-treatment with *Garcinia kola* extract and garcinoic acid reduced elevated expressions of both phospho-p65 and phospho-κBα proteins, as well as NF-κB DNA binding capacity and NF-κB-driven luciferase expression following stimulation of PBMCs with spike glycoprotein S1. Furthermore, pre-treatment of PBMCs with *Garcinia kola* extract prior to stimulation with SARS-CoV-2 spike glycoprotein S1 resulted in reduced damage to adjacent A549 lung epithelial cells. Gas Chromatography-Mass Spectrometry (GCMS) and HPLC-PDA confirmed the presence of garcinoic acid in the *Garcinia kola* extract used in this study. These results suggest that the seed of *Garcinia kola* and garcinoic acid are natural products which may possess pharmacological/therapeutic benefits in reducing cytokine storm during the late stage of severe SARS-CoV-2 and other coronavirus infections.

## 1. Introduction

Since the first report of the emergence of the severe acute respiratory syndrome coronavirus 2 (SARS-CoV-2), there has been a global spread of the infection accompanied by widespread appearance of coronavirus disease 2019 (COVID-19) (Zhu et al., 2020; Zhou et al., 2020). Among the symptoms and complications associated with SARS-CoV-2 infection, acute respiratory distress syndrome (ARDS) and organ damage have been strongly linked to disease severity and death (Moradian et al., 2020).

ARDS and multi-organ damage in SARS-CoV-2 infection have been linked to SARS-CoV-2 cytokine storm and the accompanying exaggerated release of inflammatory cytokines (Hirawat et al., 2020; Yang et al., 2020; Lee et al., 2021). In SARS-CoV-2 infection, cytokine storm involves a vicious cycle of inflammation involving excessive release of cytokines such as interleukin-1 (IL-1), interleukin-6 (IL-6), interleukin-12 (IL-12), and tumour necrosis factor (TNFα), which damage the lungs and other organs (Huang et al., 2020; Hirawat et al., 2020).

SARS-CoV-2 viral attachment and fusion to the host’s cells are facilitated by spike glycoproteins which protrude from the surface through binding to the host ACE2 receptor (Merad et al., 2020; Freeman et al., 2020). Interestingly, studies have shown that the spike glycoproteins, especially the S1 sub-unit are also a target for the host immune responses (Merad et al., 2020). Recent studies have confirmed that the S1 sub-unit induced inflammation in immune cells through activation of cellular inflammatory signalling pathways (Shirato and Kizaki, 2021; Olajide et al., 2021).

Natural products from plants have been widely suggested as potential sources of pharmacological substances for combating different components of COVID-19 pathogenesis. A molecular docking and cell-based experiments reported by Kumar et al. (2020) revealed that withanone and withaferin-A from *Withania sominifera* have the potential to interact with transmembrane protease serine 2 (TMPRSS2) and block entry of SARS-CoV-2 into cells. Also, the flavonoid narigenin was reported to be effective in effective in inhibiting human coronaviruses infection in Vero E6 cells through targeting of the endo-lysosomal two-pore channels (Clementi et al., 2021). However, there are currently no scientific report on natural products which reduce cytokine storm in COVID-19.

The edible seed of the West African plant *Garcinia kola* (bitter kola) (Figure 1A) has been widely reported to have a wide range of medicinal benefits. Extracts from *Garcinia kola* have been reported to produce anti-inflammatory activity in a number of cellular and animal models (Farombi et al., 2009; Olaleye et al., 2010; Farombi et al., 2013; Awogbindin et al., 2015; Onasanwo et al., 2016; Oyagbemi et al., 2016). Garcinoic acid (Figure 1B) is an analogue of vitamin E, which has been isolated from *Garcinia kola* seeds (Manzzini et al., 2009; Wallert et al., 2019). Garcinoic acid was reported to produce anti-inflammatory effects through inhibition of cyclo-oxygenase 2 (COX-2) and inducible nitric oxide synthase (iNOS) expressions in lipopolysaccharide (LPS)-stimulated RAW264.7 cells (Wallert et al., 2019), as well as in other cell-free and cell-based assays (Alsabil et al., 2016; Dinh et al., 2020).

**Figure 1A.**
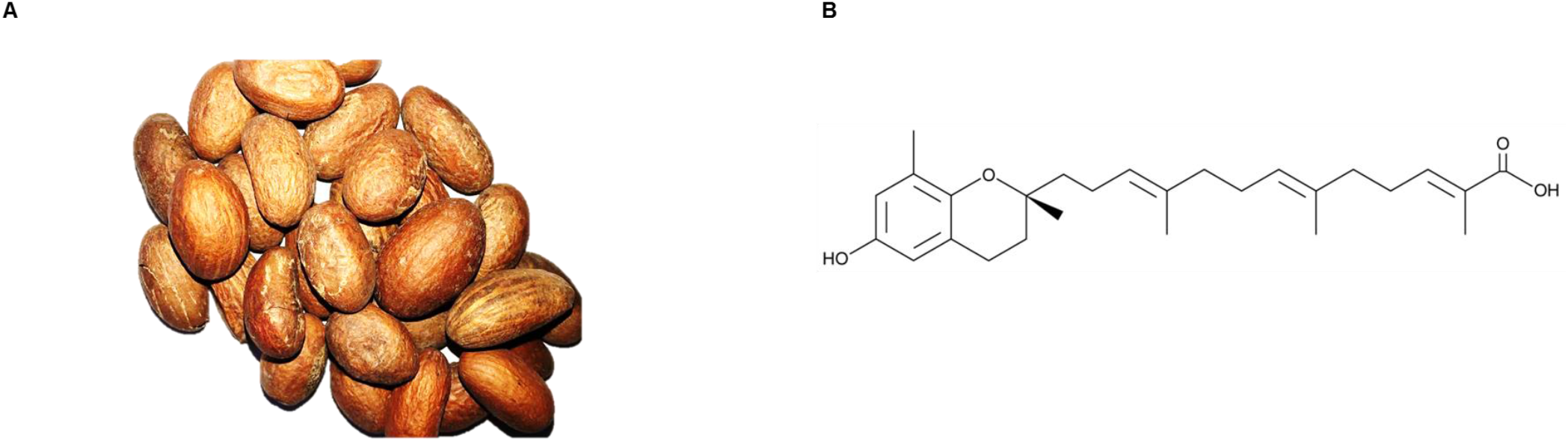
*Garcinia kola* seeds **Figure 1B**, Structure of garcinoic acid

Considering the critical role of SARS-CoV-2 cytokine storm in the pathogenesis of severe COVID-19, it is important to investigate natural products that could reduce exaggerated and organ damaging-inflammation of the disease. Consequently, we have investigated the effects of an extract of *Garcinia kola* and one of its bioactive components, garcinoic acid on SARS-CoV-2 spike glycoprotein S1-induced hyper-inflammation in PBMCs.

## 2. Materials and methods

### 2.1 Materials

Recombinant human coronavirus SARS-CoV-2 spike glycoprotein S1 (ab273068; Lots GR3356031-1 and 3353172-2; Accession MN908947) was purchased from Abcam. The protein was suspended in sterile water for functional studies. Garcinoic acid, dexamethasone and BAY11-7082 were purchased from Sigma.

### 2.2 Extraction of *Garcinia kola*

*Garcinia kola* seeds were purchased from a local herbal market in Ibadan, Nigeria and were authenticated in the herbarium of the Forestry Research Institute of Nigeria, Ibadan (Voucher Number FH1-113029). Dried seeds of *Garcinia kola* (10 g) were reduced to a fine powder in a blender and was exhaustively extracted with ethanol (300 mL). The liquid extract was filtered and dried in a petri dish at room temperature to yield the crude extract (2.6 g). This was transferred to a glass container and stored at 4°C. The extract was suspended in dimethyl sulfoxide (DMSO) for pharmacological studies. In all cases, the final concentration of DMSO in cell culture medium was 0.2%.

### 2.3 Analysis of the extract of *Garcinia kola*

Confirmation of the presence of garcinoic acid as a major component in the extract used in the assays was achieved by Gas Chromatography-Mass Spectrometry (GCMS) using pertrimethylsilyation using Pierce Tri-Sil with a characteristic molecular ion seen 571 amu (60%) (2 x tms groups) with fragmentation showing also 209 amu (100%). The extract was also analysed by HPLC-PDA using a HiChrom ACE C18 column (250mm x 4.6mm id x 3.5um) with a flow rate of 1ml/min. The linear gradient started at 90% water and 10% acetonitrile (containing 0.01% TFA) held for 4 min, rising to 100% acetonitrile over 16 minutes and held for a further 5 minutes. Garcinoic acid was observed at a retention time of 8.42 min. with UV maxima at 203 and 299 nm.

### 2.4 Cell culture

Human peripheral blood mononuclear cells (hPBMCs) (Lonza Biosciences; Catalogue #: 4W-270; Batch: 3038013) were isolated from peripheral blood by apheresis and density gradient separation. Frozen cells were thawed, and transferred to a sterile centrifuge tube. Thereafter, warmed RPMI medium was added to the cells slowly, allowing gentle mixing. The cell suspension was then centrifuged at 400 x g for 10 min. After centrifugation, the supernatant was discarded and fresh warmed RPMI was added to the pellet. This was followed by another centrifugation at 400 x g for 10 min. Supernatant was removed and cells were suspended in RPMI, counted and allowed to rest overnight. Culture plates were coated with 0.01% poly-l-lysine (Sigma) to enhance attachment of PBMCs.

### 2.5 Cell viability experiments

The MTS assay was used to assess the viability of PBMCs following treatment with either *G. kola* extract (6.25, 12.5 and 25 μg/mL) or garcinoic acid (1.25, 2.5 and 5 μM), followed by stimulation with spike protein S1 (100 ng/mL) for 24 h. At the end of the experiment, 20 μL of CellTiter 96^®^ AQueous One solution (Promega) was added to cells in a 96-well plate containing 100 μL culture medium, and incubated at 37°C for 2 h. Absorbance was read at 490nm in a Tecan Infinite M Nano microplate reader.

### 2.6 Production of pro-inflammatory cytokines

Human PBMCs were seeded out in 24-well plate at 5 × 10^4^ cells/mL and treated with *G. kola* extract (6.25, 12.5 and 25 μg/mL) or garcinoic acid (1.25, 2.5 and 5 μM) for 1 h prior to stimulation with spike protein S1 (100 ng/mL) for a further 24 h. Dexamethasone (100 nM) was used as a reference drug. Thereafter, medium was collected and centrifuged to obtain culture supernatants. Levels of TNFα in the supernatants were determined using human ELISA^™^ kit (R and D Systems). Concentrations of TNFα in supernatants were calculated from a mouse TNFα standard curve. Levels of IL-6 in supernatants were determined using human IL-6 ELISA kit (R and D Ssytems). Similarly, IL-8 production was evaluated using human IL-8 ELISA kit (R and D Systems). Absorbance measurements were carried out in a Tecan Infinite M Nano microplate reader.

### 2.7 In cell western (cytoblot) analyses

PBMCs were seeded into a 96-well plate at 5 × 10^4^ cells/mL and allowed to settle overnight. Cells were then treated with either *G. kola extract* (6.25, 12.5 and 25 μg/mL) or garcinoic acid (1.25, 2.5 and 5 μM) and incubated for 1 h prior to stimulation with spike protein S1 (100 ng/mL) for a further 15 min. At the end of each experiment, cells were fixed with 8% paraformaldehyde solution (100 μL) for 15 min., followed by washing with PBS. The cells were then incubated with primary antibodies overnight at 4°C. The following antibodies were used: rabbit anti-phospho-p65 (Cell Signalling Technology), rabbit anti-IκBα (Cell Signalling Technology), and rabbit anti-phospho-IκBα (Santa Cruz Biotechnology) antibodies. Thereafter, cells were washed with PBS and incubated with anti-rabbit HRP secondary antibody for 2 h at room temperature. Then, 100 μL HRP substrate was added to each well and absorbance measured at 450nm with a Tecan Infinite M Nano microplate reader. Readings were normalised with Janus Green normalisation stain (Abcam).

### 2.8 NF-κB p65 transcription factor binding assay

The NF-κB p65 transcription factor assay is a non-radioactive ELISA-based assay for evaluating DNA binding activity of NF-κB in nuclear extracts. PBMCs were seeded in a 6-well plate at a density of 5 × 10^4^ cells/mL. The cells were then incubated with with *G. kola extract* (6.25, 12.5 and 25 μg/mL), garcinoic acid (1.25, 2.5 and 5 μM), dexamethasone (100 nM) or BAY11-7082 (1 μM) for 1 h, followed by stimulation with spike protein S1 (100 ng/mL) for another 1 h. At the end of the incubation, nuclear extracts were prepared from the cells and subjected to NF-κB transcription factor binding assay according to the instructions of the manufacturer (Abcam).

### 2.9 Transient transfection and NF-κB reporter gene assay

Transfection of PBMCs was performed using magnetofection, a method used for transfecting primary and hard-to-transfect cells. The cells were seeded in 24-well plate at a density of 4 × 10^4^ cells/mL and allowed to settle overnight. Thereafter, culture media were replaced with Opti-MEM, with a further incubation for 2 h at 37°C. Transfection was performed by preparing a magnetofectamine O2 transfection reagent (OZ Biosciences) and Cignal NF-κB luciferase reporter (Qiagen) complex at a ratio 3:1 in 50 μL Opti-MEM. The complex was added to the 24-well plate, and the plate placed on a magnetic plate (Oz Biosciences) and incubated at 37°C for 30 min, followed by a further magnetic plate-free incubation for 18 h. Thereafter, medium was changed to culture media and cells treated with *G. kola extract* (6.25, 12.5 and 25 μg/mL), garcinoic acid (1.25, 2.5 and 5 μM), dexamethasone (100 nM) or BAY11-7082 (1 μM) for 1 h prior to stimulation with spike protein S1 (100 ng/mL) for a further 3 h. This was followed by a Dual-Glo^®^ reporter assay (Promega). Firefly and renilla luminescence intensities were measured using a FLUOstar OPTIMA microplate reader (BMG Labtech).

### 2.10 Co-culture of PBMCs with human A549 lung epithelial cells

Human A549 lung epithelial cells were co-cultured with PBMCs using the transwell system (0.4 μm porous membrane; Corning). Overnight-rested PBMCs cells were cultured in inserts that constituted the upper chamber. Thereafter, they were pre-treated with *G. kola* (6.25, 12.5 and 25 μg/mL), garcinoic acid (1.25, 2.5 and 5 μM) or dexamethasone (100 nM) for 1 h, and then incubated with spike protein S1 (100 ng/mL). One hour after the initiation of stimulation with spike protein S1, the inserts were placed on A549 cells in the lower chamber for a further 48 h. At the end of the experiment, supernatants were collected from the PBMC layer and analysed for levels of TNFα and IL-6 using human ELISA kits as described. Viability of A549 cells were also determined using the LDH assay. Briefly, 50 μL of supernatants from the A549 layer were transferred into a 96-well plate, followed by the addition of 50 μL of CytoTox 96^®^ reagent (Promega) and incubation in the dark for 30 mins at room temperature. The reaction was stopped with 50 μL stop solution and absorbance read at 490 nm.

### 2.11 Statistical analysis

Data are expressed as mean ± SEM for at least three independent experiments (n=3) and analysed using one-way analysis of variance (ANOVA) with post hoc Dunnett’s multiple comparison test. Statistical analysis were conducted using the GraphPad Prism software.

## 3. Results

### 3.1 Treatment of spike glycoprotein S1-stimulated PBMCs with extract of *Garcinia kola* and garcinoic acid did not reduce viability

PBMCs were incubated with either *Garcinia kola* extract (6.25, 12.5 and 25 μg/mL) or garcinoic acid (1.25, 2.5 and 5 μM) for 60 min and then stimulated with spike protein S1 (100 μg/mL) for a further 24 h. Results of MTS viability assays revealed that these treatments did not produce significant reduction in cell viability, in comparison with untreated cells (Figures 2A and 2B).

**Figure 2.**
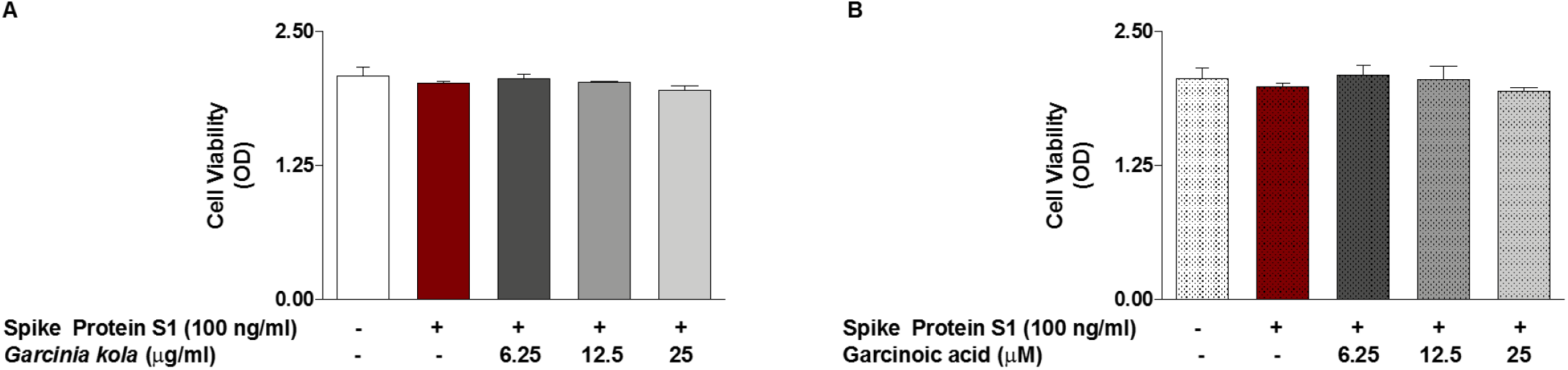
(A) MTS assay showing effects of *Garcinia kola* extract (6.25, 12.5 and 25 μg/ml) on the viability of SARS-CoV-2 spike glycoprotein S1-stimulated PBMCs. (B) MTS assay showing effects of garcinoic acid (1.25, 2.5 and 5 μM) on the viability of SARS-CoV-2 spike glycoprotein S1-stimulated PBMCs.

### 3.2 *Garcinia* kola extract reduced spike glycoprotein S1-induced increase in cytokine production

Stimulation of PBMCs with spike protein S1 (100 μg/mL) for 24 h resulted in significant (p<0.001) increase in the release of TNFα into culture supernatants, when compared with unstimulated cells. However, pre-treatment with *G. kola* (6.25, 12.5 and 25 μg/mL) for 60 min prior to spike protein S1 stimulation resulted in significant (p<0.05) and concentration-dependent reduction in TNFα production (Figure 3A). The highest concentration of *G. kola* investigated (25 μg/mL) reduced spike protein S1-induced increased TNFα production by ~47%, while dexamethasone (100 nM) treatment caused a reduction by ~75% (Figure 3A). Similarly, spike protein S1-induced increased release of IL-6 was significantly reduced (p<0.01) when cells were pre-treated with *G. kola* (12.5 and 25 μg/mL), and dexamethasone (Figure 3B). Analyses of culture supernatants further revealed that spike protein S1 stimulation of PBMCs resulted in significant elevation of IL-1β (Figure 3C) and IL-8 (Figure 3D), which were reduced in the presence of *G. kola* extract (6.25-25 μg/mL).

**Figure 3.**
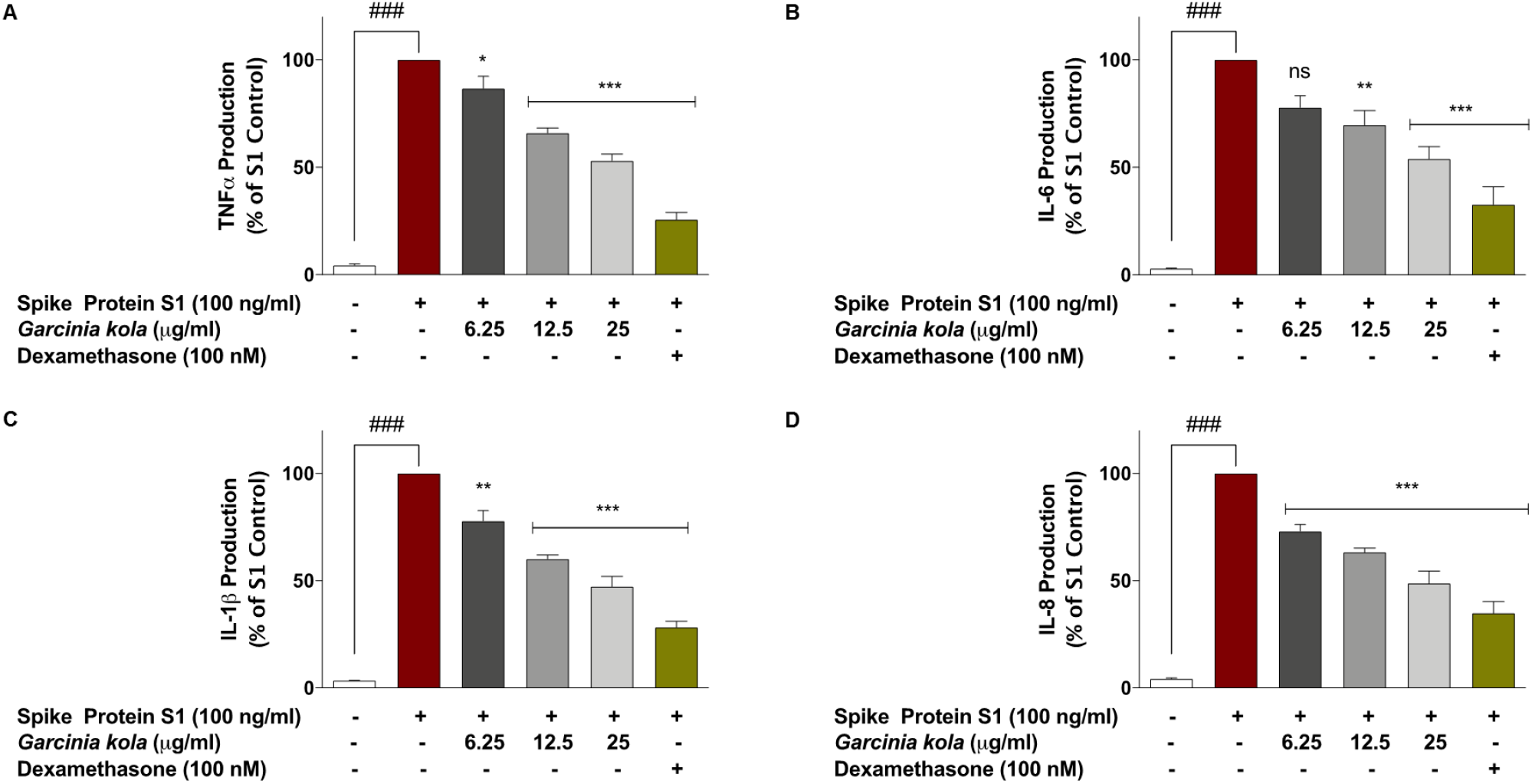
Pre-treatment of SARS-CoV-2 spike glycoprotein S1-stimulated PBMCs with *Garcinia kola* extract (6.25, 12.5 and 25 μg/ml) and dexamethasone (100 nM) reduced increased production of TNFα (A), IL-6 (B), IL-1β (C), and IL-8 (D). Levels of cytokines in cell supernatants were detected using human TNFα, IL-6, IL-1β, and IL-8 ELISA kits. Values are mean ± SEM for at least 3 independent experiments (ns: not significant; ### p<0.001 unstimulated control *versus* SARS-CoV-2 spike glycoprotein S1 stimulation. *p<0.05; **p<0.01; ***p<0.001, treatments *versus* SARS-CoV-2 spike glycoprotein S1 stimulation; one-way ANOVA with post-hoc Dunnett’s multiple comparison test).

### 3.3 Effects of garcinoic acid pre-treatment on spike glycoprotein S1-induced increased production of pro-inflammatory cytokines

Experiments on garcinoic acid showed that spike protein S1-induced increased production of TNFα was significantly reduced by ~22%, 35% and 53% in cells pre-treated with 1.25, 2.5 and 5 μM of the compound, respectively (Figure 4A). S1-induced increased production of IL-6, IL-1β and IL-8 were also significantly (p<0.01) reduced when cells were pre-treated with 1.25, 2.5 and 5 μM of garcinoic acid (Figures 4B, 4C and 4D). At 5 μM, garcinoic acid reduced increased production of IL-6, IL-1β and IL-8 by ~45%, ~53% and ~52%, respectively. However, dexamethasone (100 nM) reduced their levels by ~67%, ~75% and ~70%, respectively.

**Figure 4.**
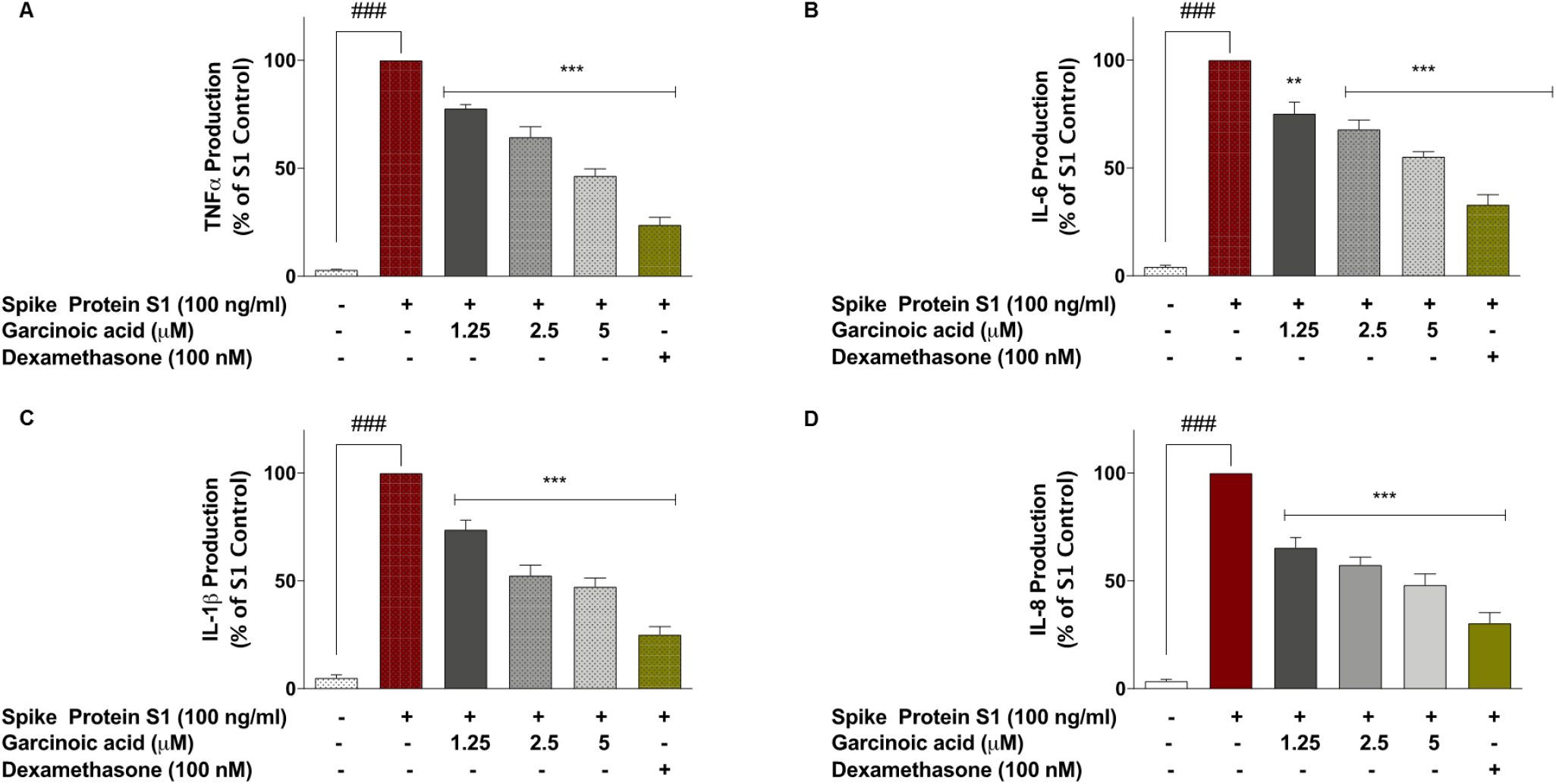
Pre-treatment of SARS-CoV-2 spike glycoprotein S1-stimulated PBMCs with garcinoic acid (1.25, 2.5 and 5 μM) and dexamethasone (100 nM) reduced increased production of TNFα (A), IL-6 (B), IL-1β (C), and IL-8 (D). Levels of cytokines in cell supernatants were detected using human TNFα, IL-6, IL-1β, and IL-8 ELISA kits. Values are mean ± SEM for at least 3 independent experiments (### p<0.001 unstimulated control *versus* SARS-CoV-2 spike glycoprotein S1 stimulation. **p<0.01; ***p<0.001, treatments *versus* SARS-CoV-2 spike glycoprotein S1 stimulation; one-way ANOVA with post-hoc Dunnett’s multiple comparison test).

### 3.4 *Garcinia kola* extract and garcinoic acid inhibit spike glycoprotein S1-induced inflammation by targeting NF-κB activation

Encouraged by results showing reduction in spike protein S1-induced increase in pro-inflammatory cytokine production by both *G. kola* extract and garcinoic acid, experiments were conducted to determine whether inhibition of NF-κB activation contributed to their effects. Stimulation of PBMCs with (100 ng/mL) spike protein S1 resulted in ~8.8-fold and ~8.2-fold increases in protein expressions of phoshpho-p65, and phospho-IκBα, respectively (Figures 5A and 5B). However, pre-treating the cells with *G. kola* extract (6.25, 12.5 and 25 μg/mL) for 60 min prior to spike protein S1 stimulation resulted in significant (p<0.01) reduction in protein levels of both phospho-p65 and phospho-IκBα (Figures 5A and 5B). Also, expressions of both proteins were significantly reduced in cells pre-treated with dexamethasone (100 nM) and the NF-κB inhibitor, BAY11-7082 (1 μM), in comparison with cells stimulated with spike protein S1 alone.

**Figure 5.**
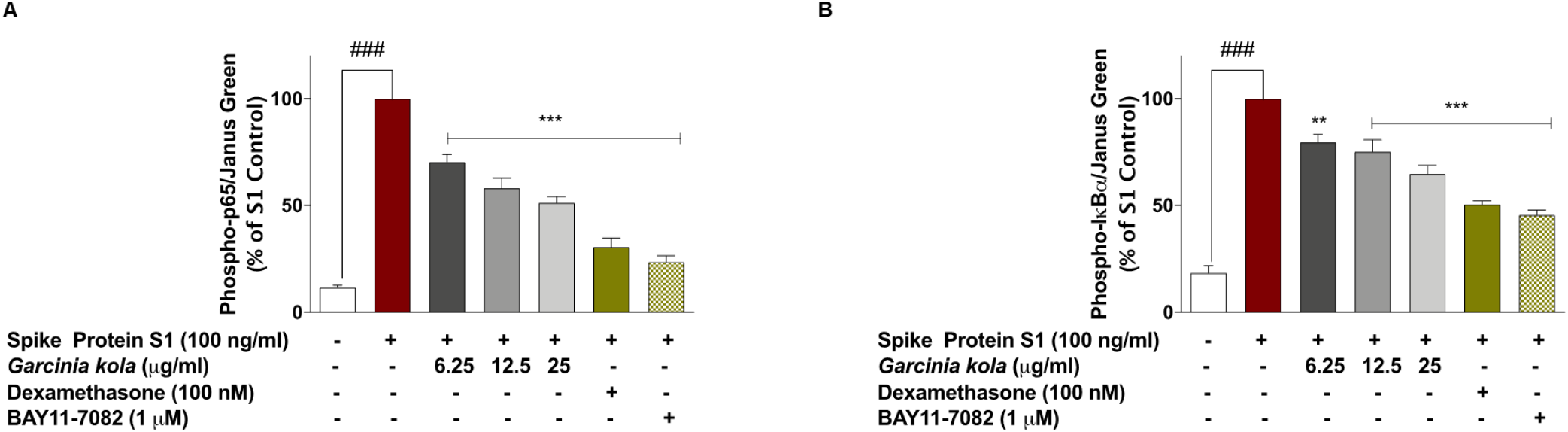
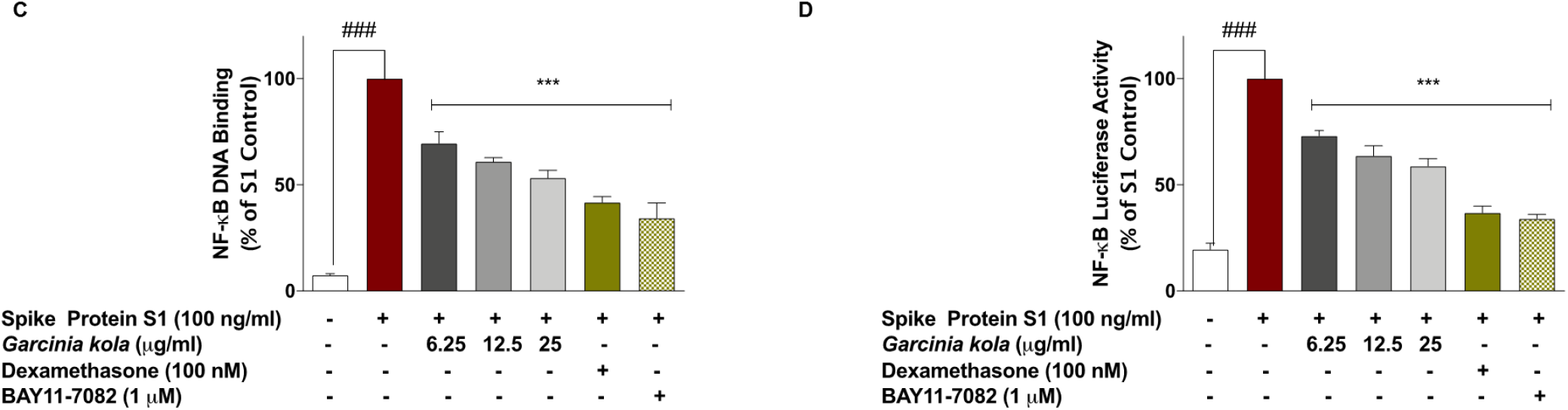
SARS-CoV-2 spike glycoprotein S1-induced increased expression of phospho-p65 (A), phospho-IκBα (B) and NF-κB p65 DNA binding (C) in PBMCs was inhibited in the presence of *Garcinia kola* extract (6.25, 12.5 and 25 μg/mL), dexamethasone (100 nM) and BAY11-7082 (1 μM). Cells were stimulated for 15 min, followed by in cell western analysis with anti-phospho-p65 and anti-phospho-IκBα antibodies. DNA binding was evaluated using NF-κB transcription factor binding assay kit. (D) Effects of *Garcinia kola* extract (6.25, 12.5 and 25 μg/mL), dexamethasone (100 nM) and BAY11-7082 (1 μM) on NF-κB-driven luciferase activity in SARS-CoV-2 spike glycoprotein S1-stimulated PBMCs. Values are mean ± SEM for at least 3 independent experiments (### p<0.001 unstimulated control *versus* SARS-CoV-2 spike glycoprotein S1 stimulation. **p<0.01; ***p<0.001, treatments *versus* SARS-CoV-2 spike glycoprotein S1 stimulation; one-way ANOVA with post-hoc Dunnett’s multiple comparison test).

Further experiments to evaluate effects of *G. kola* extract on spike protein S1-induced increased activation of NF-κB signalling revealed that in the presence of 6.25, 12.5 and 25 μg/mL of the extract, DNA binding capacity of p65 sub-unit was reduced by ~31%, ~39% and ~47%, respectively when compared with S1-stimulated PBMCs. Results also showed that spike protein S1-induced DNA binding capacity was reduced by ~58% and ~66% when PBMCs were pre-treated with dexamethasone (100 nM) and BAY11-7082 (1 μM), respectively (Figure 5C). Results of reporter gene assays in Figure 5D show that stimulation of NF-κB luciferase reporter transfected cells with spike protein S1 resulted in activation of NF-κB-driven luciferase expression, which was significantly (p<0.001) reduced by *G. kola* extract (6.25, 12.5 and 25 μg/mL), dexamethasone (100 nM) and BAY11-7082 (1 μM).

Investigations to determine the role of NF-κB inhibition in the effects of garcinoic acid on spike protein S1-induced hyper-inflammation in PBMCs revealed that the compound produced significant (p<0.01) inhibition of p65 and IκBα phosphorylation (Figures 6A and 6B). Experiments also showed that garcinoic acid (5 μM) inhibited spike protein S1-induced NF-κB DNA binding capacity and NF-κB-driven luciferase expression by ~51% and ~42%, respectively. Dexamethasone (100 nM) reduced these events by ~64% and ~61%, respectively (Figures 6C and 6D).

**Figure 6.**
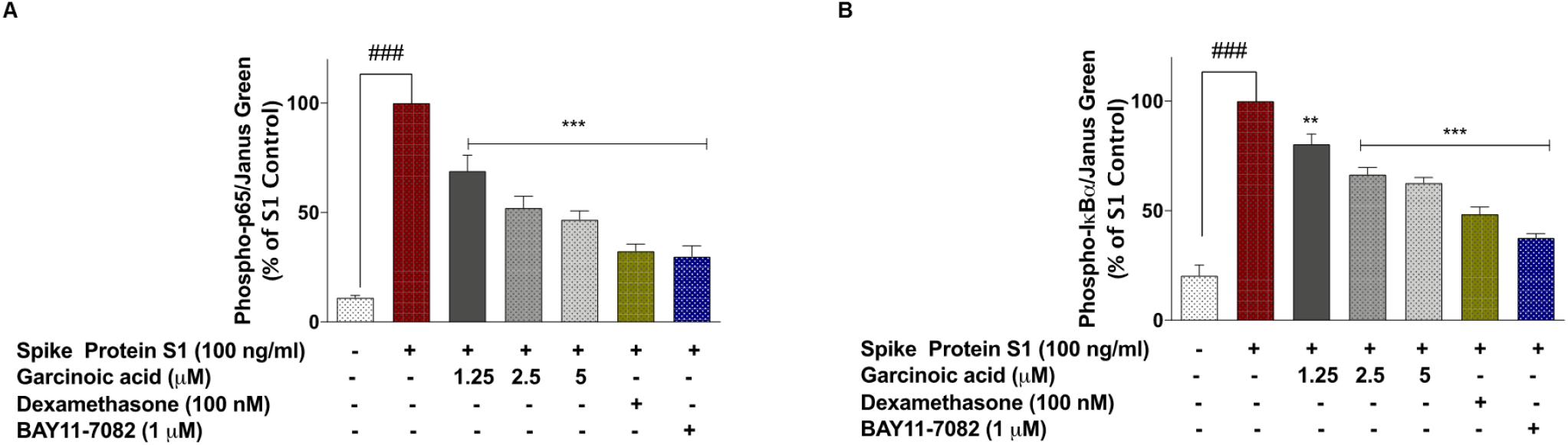
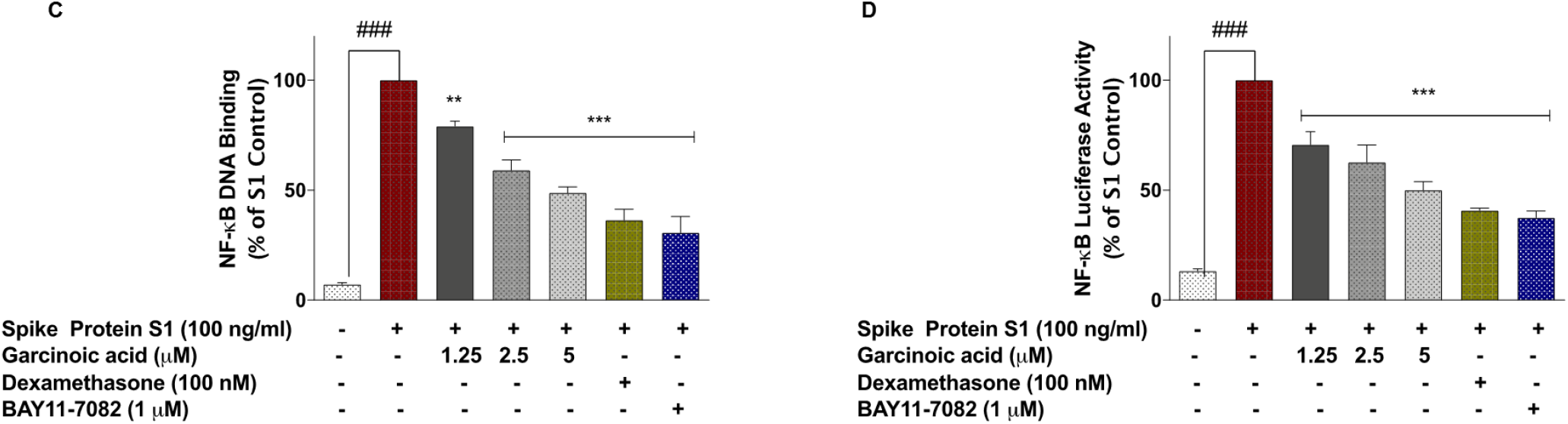
SARS-CoV-2 spike glycoprotein S1-induced increased expression of phospho-p65 (A), phospho-IκBα (B) and NF-κB p65 DNA binding (C) in PBMCs was inhibited in the presence of garcinoic acid (1.25, 2.5 and 5 μM), dexamethasone (100 nM) and BAY11-7082 (1 μM). Cells were stimulated for 15 min, followed by in cell western analysis with anti-phospho-p65 and anti-phospho-IκBα antibodies. DNA binding was evaluated using NF-κB transcription factor binding assay kit. (D) Effects of garcinoic acid (1.25, 2.5 and 5 μM), dexamethasone (100 nM) and BAY11-7082 (1 μM) on NF-κB-driven luciferase activity in SARS-CoV-2 spike glycoprotein S1-stimulated PBMCs. Values are mean ± SEM for at least 3 independent experiments (### p<0.001 unstimulated control *versus* SARS-CoV-2 spike glycoprotein S1 stimulation. **p<0.01; ***p<0.001, treatments *versus* SARS-CoV-2 spike glycoprotein S1 stimulation; one-way ANOVA with post-hoc Dunnett’s multiple comparison test).

### 3.3 Effects of *Garcinia kola* extract and garcinoic acid on the viability of A549 human lung alveolar epithelial cells co-cultured with spike glycoprotein S1-stimulated PBMCs

As shown in Fgiures 7A and 8A, stimulation of PBMCs with spike glycoprotein S1 (100 ng/mL) for 48 h resulted in a significant (p<0.001) increase in the release of LDH by adjacent A549 lung epithelial cells. This was accompanied by significant increases in the levels of both TNFα (Figures 7B and 8B) and IL-6 (Figures 7C and 8C) production by the PBMCs. Pre-treatment of PBMCs with 6.25 μg/mL of *G. kola* extract prior to stimulation with spike glycoprotein S1 did not result in significant (p<0.05) reduction in cytotoxicity to co-cultured A549 lung epithelial cells. However, on increasing the concentrations of *G. kola* extract to 12.5 and 25 μg/mL, significant (p<0.05) reduction in damage to A549 cells was observed (Figure 7A). Interestingly, levels of TNFα and IL-6 were significantly (p<0.001) reduced in the presence of all concentrations of the extract tested (Figures 7B and 7C).

**Figure 7.**
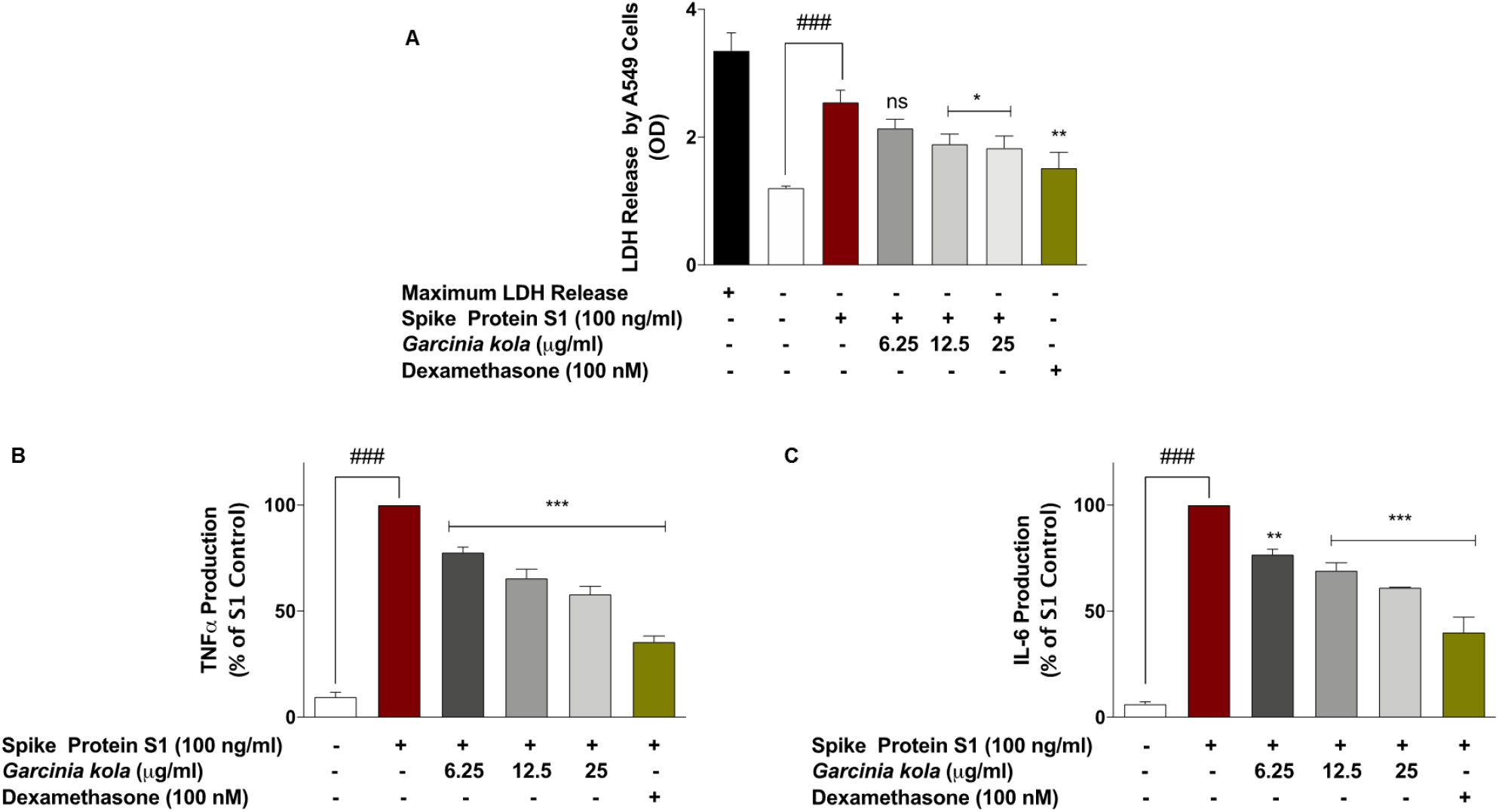
*Garcinia kola* extract (12.5 and 25 μg/mL) and dexamethasone (100 nM) prevented reduced viability of A549 lung epithelial cells co-cultured with SARS-CoV-2 spike glycoprotein S1-stimulated PBMCs (A), accompanied by reduction in TNFα and IL-6 levels in the PBMC layer of the co-culture (C and D). Values are mean ± SEM for at least 3 independent experiments (ns: not significant; ### p<0.001 unstimulated control *versus* SARS-CoV-2 spike glycoprotein S1 stimulation. *p<0.05; **p<0.01; ***p<0.001, treatments *versus* SARS-CoV-2 spike glycoprotein S1 stimulation; one-way ANOVA with post-hoc Dunnett’s multiple comparison test).

**Figure 8.**
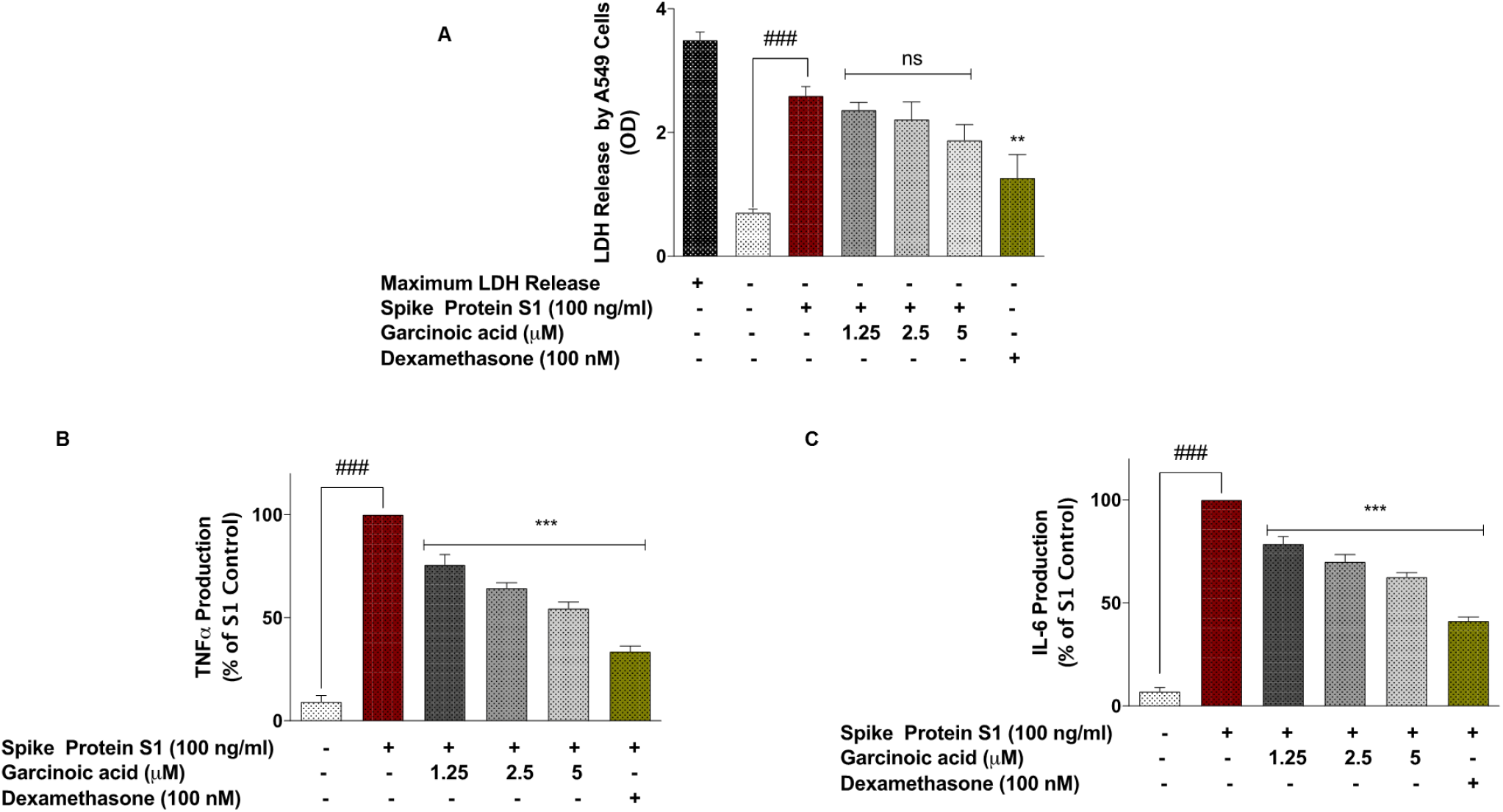
Dexamethasone (100 nM), but not garcinoic acid (1.25, 2.5 and 5 μM) significantly prevented reduced viability of A549 lung epithelial cells co-cultured with SARS-CoV-2 spike glycoprotein S1-stimulated PBMCs (A), accompanied by reduction in TNFα and IL-6 levels in the PBMC layer of the co-culture (C and D). Values are mean ± SEM for at least 3 independent experiments (ns: not significant; ### p<0.001 unstimulated control *versus* SARS-CoV-2 spike glycoprotein S1 stimulation. **p<0.01; ***p<0.001, treatments *versus* SARS-CoV-2 spike glycoprotein S1 stimulation; one-way ANOVA with post-hoc Dunnett’s multiple comparison test).

Pre-treatment of spike glycoprotein S1-stimulated PBMCs with garcinoic acid (1.25, 2.5 and 5 μM) did not significantly (p<0.05) prevent toxicity to adjacent A549 cells after 48 h (Figure 8A). However, there were significant (p<0.001) reductions in the levels of both TNFα and IL-6 in the PBMC layer of the co-culture in the presence of all the concentrations of the compound (Figures 8B and 8C).

## 4. Discussion

Therapeutic strategies have been proposed for COVID-19, the disease caused by infection with the SARS-CoV-2 virus. Among these are potential treatments for COVID-19 cytokine storm (Tang et al., 2020). In this regard, anti-inflammatory steroids such as dexamethasone have been shown to provide some benefits in the treatment of COVID-19, based on results of RECOVERY and CoDEX trials (RECOVERY Collaborative Group, 2021; Tomazini et al., 2020; Ledford, 2020). Dexamethasone and other corticosteroids were also reported to provide benefits in previous coronavirus outbreaks, such as Middle East respiratory syndrome (MERS) and severe acute respiratory syndrome (SARS) (Stockman et al., 2006; Chen et al., 2006; Arabi et al., 2018). The benefits of dexamethasone and other corticosteroids in the treatment of coronavirus infections are linked to their anti-inflammatory activity in suppressing cytokine storms. In spite of the potential benefits of dexamethasone in the treatment of severe COVID-19 symptoms, concerns have been raised about enhancement of viral replication due to immunosuppressive property of the drug. Therefore, the use of alternative anti-inflammatory modalities, including natural products could provide an alternative therapeutic strategy in treating COVID-19 cytokine storm.

This study showed that an extract from *Garcinia kola* seeds reduced the production of pro-inflammatory cytokines TNFα, IL-6, IL-1β and IL-8 in in human PBMCs stimulated with SARS-CoV-2 spike protein S1. These are significant observations as clinical evidence has shown that the cytokine storm in COVID-19 involves a vicious cycle of inflammatory responses which are characterised by excessive release of IL-1β, IL-6, TNFα, which then target lung and other tissues (Hirawat et al., 2020; Grifoni et al., 2020; Aziz et al., 2020). The effects of *Garcinia kola* extract against SARS-CoV-2 spike protein S1-induced increased production of pro-inflammatory cytokines in PBMCs are consistent with previous reports of similar anti-inflammatory activities *in vivo* and *in vitro* (Farombi et al., 2009; Olaleye et al., 2010; Farombi et al., 2013; Onasanwo et al., 2016; Oyagbemi et al., 2016). Furthermore, a study reported by Awogbindin et al. (2015), shows that a seed extract of *Garcinia kola* attenuated inflammatory cell infiltration in a mouse model of influenza A virus.

Garcinoic acid is a δ-tocotrienol derivative, and one of the major bioactive components in the seeds of *Garcinia kola.* Similar to activities shown by the seed extract of *Garcinia kola,* garcinoic acid significantly inhibited SARS-CoV-2 spike protein S1-induced exaggerated production of TNFα, IL-6, IL-1β and IL-8 in PBMCs. It therefore appears that garcinoic acid may be contributing to the anti-inflammatory effects shown by *Garcinia kola* extract, as the compound was previously shown to produce anti-inflammatory effects in LPS-stimulated RAW264.7 cells (Wallert et al., 2019). Further phytochemical analyses are required to identify other bioactive components of *Garcinia kola* with similar pharmacological profiles.

Activation of the NF-κB transcription factor signalling pathway has been linked to the pathogenesis of severe COVID-19 (Hirano and Murakami 2020). Furthermore, a review by Hariharan et al. (2021) suggests that inhibition of NF-κB activation, with the resultant reduction in exaggerated production of pro-inflammatory cytokines like TNFα will potentially result in a reduction in cytokine storm in severe COVID-19. Our recent studies also showed that NF-κB activation is a mechanism for SARS-CoV-2 spike protein S1-induced exaggerated production of TNFα, IL-6, IL-1β and IL-8 in PBMCs (Olajide et al., 2021). The seed extract of *Garcinia kola* and garcinoic acid inhibited both cytoplasmic activation, DNA binding and transcriptional activity of NF-κB in SARS-CoV-2 spike protein S1-simulated PBMCs, suggesting a role for the transcription factor in their anti-inflammatory activities. The effects of *Garcinia kola* on NF-κB activation in this study appear to be similar to previously observed activities in RAW264 macrophages and BV-2 microglia stimulated with bacterial lipopolysaccharide (Abarikwu, 2016; Onasanwo et al., 2016). Similarly, inhibition of NF-κB activation in LPS-stimulated macrophages has been reported for garcinoic acid (Schmölz et al., 2014). Further investigations are required to identify specific molecular targets involved in NF-κB-mediated inhibition of SARS-CoV-2 spike protein S1-induced inflammation by *Garcinia kola* and garcinoic acid.

Several studies have implicated COVID-19 cytokine storm in lung damage and ARDS (Wang et al., 2020). In this regard, SARS-CoV-2 is known to activate immune cells leading to the leads to the production of a large number of pro-inflammatory cytokines which cause lung damage. Results of this study showed that *Garcinia kola* seed extract but not garcinoic acid prevented toxicity to A549 lung epithelial cells which were co-cultured with SARS-CoV-2 spike protein S1-stimulated PBMCs, suggesting that the anti-inflammatory activity of *Garcinia kola* resulted in the prevention of cytokine-mediated damage to A549 lung epithelial cells. It is proposed that the anti-inflammatory effects of *Garcinia kola* seed extract could be an important pharmacological attribute for the treatment of ARDS and lung damage in patients with severe COVID-19.

Anti-inflammatory strategies have shown promising results in the alleviation of ARDS and lung damage resulting from COVID-19. Tocilizumab (a potential recombinant monoclonal antibody against IL-6) is currently under investigation for the management of ARDS in patients with COVID-19 (Khiali et al., 2020; Nasir et al., 2020; Toniati et al., 2020). Similarly anti-inflammatory drugs bazedoxifene and raloxifene, used in arthritis have been proposed as treatments for preventing cytokine storm, ARDS and mortality in severe COVID-19 patients (Smetana et al., 2020).

While these results are based on the outcome of *in vitro* experiments on human PBMCs, they do suggest that the seed of *Garcinia kola* and garcinoic acid are natural products which may possess pharmacological/therapeutic benefits in reducing cytokine storm during the late stage of severe SARS-CoV-2 and other coronavirus infections.

## Acknowledgements

We thank Tobi Olajide for providing technical assistance with the processing of *Garcinia kola* seeds.

## Notes

### Competing Interest Statement

The authors have declared no competing interest.

